# Evaluation of Host Plant Preferred by Asian Citrus Psyllid – *Diaphorina citri*

**DOI:** 10.1101/2025.03.04.641519

**Authors:** Umang Deep Rai, Jigme Thinley

**Affiliations:** Master of Science in Agriculture and Food Security, Edith Cowan University, Western Australia; Master of Food Security, Murdoch University, Western Australia

**Keywords:** citrus, *Diaphorina citri*, host-preference, Huanglongbing, oviposition

## Abstract

Huanglongbing, also known as citrus greening, is a serious disease affecting citrus crops in Bhutan, leading to a decline in yield. The Asian citrus psyllid (*Diaphorina citri*) serves as the vector for this disease. A Completely Randomized Design (CRD) experiment was conducted with four treatments and three replications to assess the effectiveness of various alternate host plants in attracting the psyllid. The experiment involved a choice test with four host plants: Murraya paniculata (orange jessamine), Murraya koenigii (curry leaf plant), Citrus aurantifolia (Himalayan lime), and Citrus reticulata (mandarin). For each treatment, the numbers of eggs, nymphs, and adults were recorded. Data analysis was performed using One-way ANOVA, with significance set at P < .05. It was confirmed that the Asian citrus psyllid oviposits on all four host plants. Notably, the psyllid completed its life cycle the fastest on the curry leaf plant, taking an average of 13.33 days. The highest nymphal mortality was observed in mandarin and Himalayan lime at 41%, whereas the lowest mortality rate occurred when all four host plants were present, at 21%. The curry leaf plant was identified as the most effective at attracting the psyllid, followed by orange jessamine. These two species could be utilized as trap crops for psyllid management in orchard settings.

## Introduction

Citrus cultivation is a crucial agricultural sector in Bhutan, contributing significantly to the economy and nutrition of its population. This fruit is cultivated across 17 Dzongkhags (districts) and is recognized for its economic value. However, citrus trees are susceptible to Huanglongbing disease, which is transmitted by the Asian citrus psyllid (Diaphorina citri). The Asian citrus psyllid is regarded as the most critical pest affecting citrus, as it acts as a vector for the bacterium Candidatus Liberibacter asiaticus, the causative agent of Huanglongbing [4]. This disease is responsible for the deterioration of citrus orchards in Bhutan and can lead to the complete loss of citrus trees[10].

Huanglongbing, commonly referred to as citrus greening, poses a significant threat to citrus yields globally, affecting various regions including China, the Philippines, the Arabian Peninsula, and parts of Africa [1]. It is classified as one of the most destructive and challenging diseases for citrus farmers, resulting in substantial yield declines. The crisis necessitates urgent attention to revive the citrus industry in Bhutan[10]. The presence of Huanglongbing in Bhutanese orchards was definitively confirmed in 2002 in the Punakha and Wangduephodrang Dzongkhags, along with the identification of its vector. Symptoms linked to Huanglongbing and its vectors were first noted in the mid-1990s [3]. The disease is thought to be a primary factor contributing to citrus decline in Bhutan, particularly in low-lying regions situated below 1,000 meters above sea level [20].

While there are established chemical methods for controlling psyllids [15] there is a pressing need for further research to develop alternative management strategies. One approach is the use of alternative host plants as trap crops within citrus orchards. Identifying these alternative host plants is crucial for the effective management of Huanglongbing. Therefore, this study aims to assess the alternative host plants preferred by D. citri under controlled laboratory conditions.

## Methods and Materials

### Site/study area

The experimental research was carried out at the Agriculture Research and Development Centre (ARDC) in Bajo, Wangduephodrang Dzongkhag. The laboratory experiment took place from January to April 2020, specifically in the Plant Protection laboratory. Located approximately 69 km east of Thimphu, the site sits at an elevation of around 1200 meters above sea level [11].

### Materials used

This experimental study consisted of a choice experiment featuring four distinct treatments. The treatments included: T1 – mandarin and Himalayan lime; T2 – mandarin and curry leaf plant; T3 – mandarin and orange jessamine; and T4 – a combination of mandarin, Himalayan lime, curry leaf plant, and orange jessamine.

### Experimental set-up

For the collection and rearing of psyllids, we utilized an insect-rearing cage, manual aspirators, and tapping trays. A data logger was implemented to monitor daily temperature and relative humidity levels in the laboratory. Asian citrus psyllids were collected using manual aspirators[7]. To ensure adequate humidity in the lab, muslin cloths soaked in water were placed in bowls. The host plants were cultivated in polyethylene pots filled with a 1:1 mixture of soil and leaf mold.

Asian citrus psyllid was collected from citrus trees in the Kamichhu Royal Orchard in Wangduephodrang Dzongkhag. Bugdorm insect cages of 45 cm x 45 cm x 45 cm were used for the transportation of Asian citrus psyllids from the orchard to the laboratory.

### Data collection

Eggs, adults, and nymphs were counted daily throughout the life cycle of the psyllids. The total counts for each category—eggs, nymphs, and adults—were meticulously recorded for each treatment. Data collection occurred each day at 10:00 a.m. The overall duration of the life cycle on the host plants was documented, as referenced in sources [8] and [18]. The parameters, including the counts of eggs, nymphs, and adults, were systematically organized into tables for all four treatments. Data collection was carried out individually for each treatment.

### Data analysis

The software utilized for data recording was Microsoft Excel 2016[9]. The experimental data across the four treatments were analyzed using appropriate Analysis of Variance (ANOVA). Statistix 8 was employed to perform a one-way ANOVA within a Completely Randomized Design, enabling us to evaluate significant differences in the mean counts of eggs, nymphs, and adults on the host plants for each treatment. The counts of eggs, nymphs, and adults for each treatment were sampled and averaged as needed. The differences in treatment means were considered statistically significant at *P =* .*05* level. The rate of nymphal mortality was calculated in percentage.

## Results

### Oviposition

The introduced adult Asian citrus psyllids laid their eggs on the new flush shoots of all four host plants used in the experiment. Oviposition occurred 3 to 5 days after the introduction of the adult psyllids into the treatments. The presence of young flush significantly influenced egg-laying on the host plants, as reported by sources [18, 19, 16]. The eggs, which were yellowish in color, were found on the upper surfaces of the leaves.

### Incubation

The incubation period for the Asian citrus psyllids was 4–6 days in all the treatments. This is in line with the findings by [18], stating that at 25°C, psyllid eggs hatch in about 4 days.

### Nymph and adult stages

Nymphs were difficult to observe without the help of a magnifying glass. Both nymphs and adults had a waxy secretion, although more secretion occurred after the appearance of nymphs. Nymphs were greenish to yellow in colour. They were found to be docile. Adults were visible without using magnifying glass. The adults were found at 45 degree angle to the shoot or leaf upon which they stood. Adults were dark coloured.

### Choice experiment

#### Mandarin and Himalayan lime

The results of the study comparing preference between the host plants—mandarin and Himalayan lime—show a significant difference (p < .05) in the number of eggs deposited on each plant (refer to Table 1). The mandarin exhibited a higher quantity of eggs (13.67 ± 0.67), nymphs (12.67 ± 1.45), and adults (5.33 ± 0.67). These findings indicate that Himalayan lime is not an effective host plant for the Asian citrus psyllid.

**Table 1.** Total number of eggs, nymphs and adult *D. citri* present on choice situation (Mandarin and Himalayan lime).

| Parameters | Mandarin | Himalayan lime | CV% | F test |
| --- | --- | --- | --- | --- |
| Number of eggs | 13.67 ± 0.67<sup>a</sup> | 5.7 ± 1.15<sup>b</sup> | 17.50 | ** |
| Number of nymphs | 12.67 ± 1.45<sup>a</sup> | 4.37 ± 0.67<sup>b</sup> | 23.03 | ** |
| Number of adults | 5.33 ± 0.67<sup>a</sup> | 1.67 ± 0.89<sup>b</sup> | 38.69 | ** |

Means with different superscript within the rows differs significantly ( $P < .05$ ), \*\*\* = significant difference at $p < .001$ , \*\* = significant difference at $p < .05$ , CV = coefficient of variation.

### Mandarin and curry leaf plant

The comparison of preferences between the two host plants, mandarin and curry leaf, reveals a significant difference (p < 0.05) in the total counts of eggs, nymphs, and adults (refer to Table 2). The curry leaf plant recorded the highest average number of eggs at 17.67 ± 0.33, while the mandarin averaged 10.33 ± 0.88. This indicates that Asian citrus psyllids exhibit a preference for the curry leaf plant over the mandarin for oviposition and breeding.

**Table 2.** Total number of eggs, nymphs and adult *D. citri* present on choice situation (Mandarin and Curry leaf plant).

| Parameters | Mandarin | Curry leaf plant | CV% | F test |
| --- | --- | --- | --- | --- |
| Number of eggs | 10.33 ± 0.88<sup>b</sup> | 17.67 ± 0.33<sup>a</sup> | 8.25 | ** |
| Number of nymphs | 4.67 ± 0.33<sup>b</sup> | 21 ± 0.57<sup>a</sup> | 6.36 | *** |
| Number of adults | 2 ± 0.57<sup>b</sup> | 16.67 ± 0.67<sup>a</sup> | 11.57 | *** |

Means with different superscript within the rows differs significantly ( $P < .05$ ), \*\*\* = significant difference at $p < .001$ , \*\* = significant difference at $p < .05$ , CV = coefficient of variation.

### Mandarin and orange jessamine

The findings regarding the preference between the two host plants, mandarin and orange jessamine, reveal a significant difference (P < .05) in the total quantities of eggs, nymphs, and adults (as shown in Table 3). Orange jessamine exhibited higher averages, with 18.33 ± 0.33 eggs, 16.67 ± 0.33 nymphs, and 12.67 ± 0.33 adults, in comparison to mandarin.

**Table 3.** Total number of eggs, nymphs and adult *D. citri* present on choice situation (Mandarin and Orange jessamine).

| Parameters | Mandarin | Orange jessamine | CV | F test |
| --- | --- | --- | --- | --- |
| Number of eggs | 11.67 ± 0.88<sup>b</sup> | 18.33 ± 0.33<sup>a</sup> | 7.70 | ** |
| Number of nymphs | 5.67 ± 0.33<sup>b</sup> | 16.67 ± 0.33<sup>a</sup> | 5.17 | *** |
| Number of adults | 1.33 ± 0.33<sup>b</sup> | 12.67 ± 0.33<sup>a</sup> | 8.25 | *** |

Means with different superscript within the rows differs significantly ( $P < .05$ ), \*\*\* = significant difference at $p < .001$ , \*\* = significant difference at $p < .05$ , CV = coefficient of variation.

### Mandarin, orange jessamine, curry leaf plant and Himalayan lime

In Treatment 4, a significant difference (P < .05) was found in the number of eggs, nymphs, and adults laid by D. citri among the four host plants (refer to Table 4). The Bonferroni multiple comparison test revealed that the curry leaf plant had the highest average number of eggs (22 ± 0.33), while the Himalayan lime exhibited the lowest average (6.67 ± 0.88), as illustrated in Table 4. Furthermore, the curry leaf plant also supported the highest number of adults (17.67 ± 0.33), whereas the Himalayan lime showed no adults present (0.00 ± 0.57). While oviposition occurred on all four host plants, higher survival rates for nymphs and adults were observed on the curry leaf plant and orange jessamine compared to mandarin and Himalayan lime. These findings indicate that the curry leaf plant is the preferred host for oviposition, as well as for nymphal and adult development.

**Table 4.**
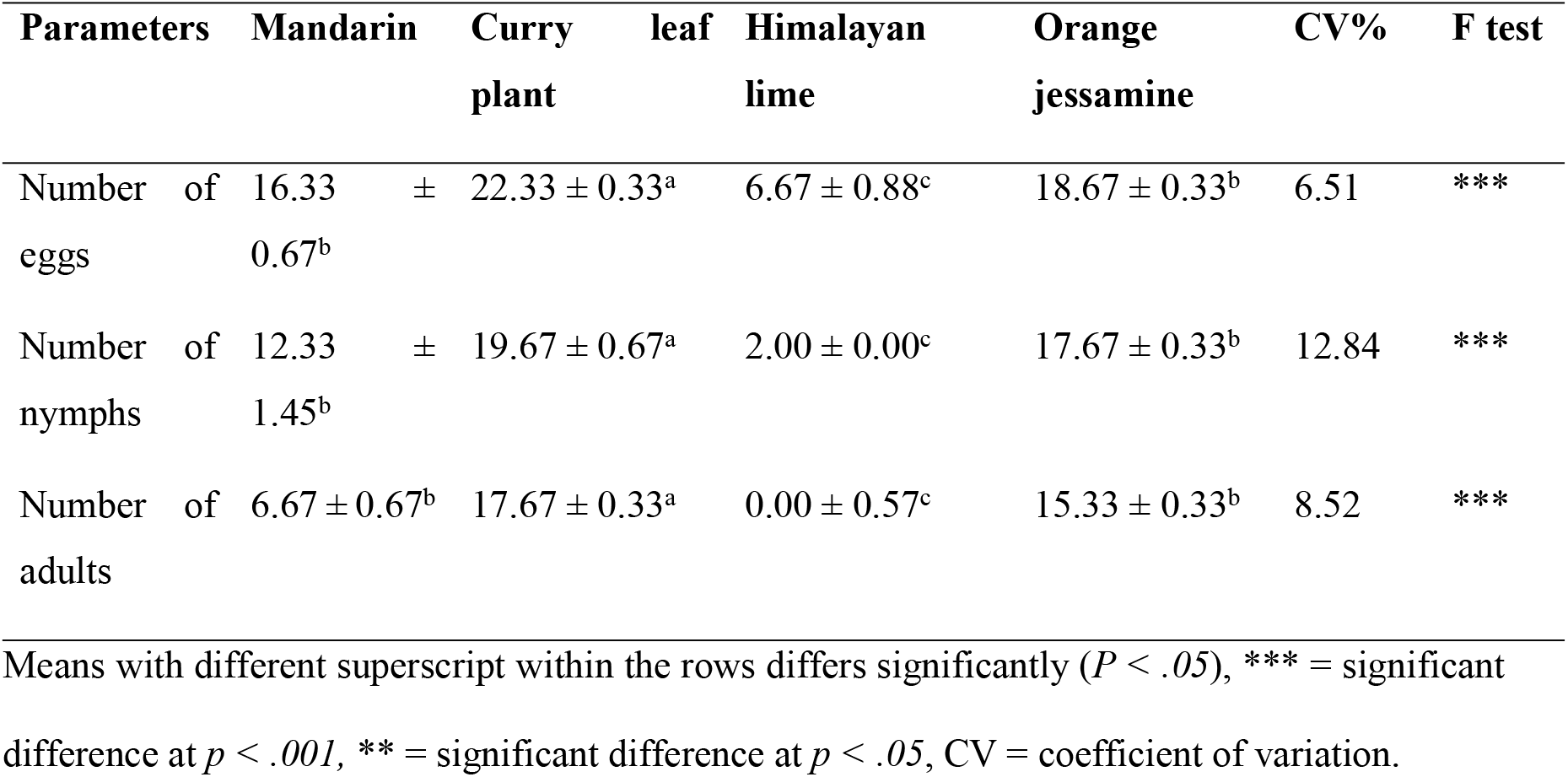
Total number of eggs, nymphs and adult D. citri present on four choice host plants.

### Comparative life cycle of *D. citri*

The mean incubation period for the eggs did not show a significant difference (P > .05), with an average of 5.67 days observed for the Himalayan lime and a shorter average of 3.33 days for the curry leaf plant (see Table 5). The overall developmental period from egg to adult stages took 22.00 days on the Himalayan lime, while it was reduced to 16.66 days on the curry leaf plant.

**Table 5.** Egg hatching and development time (days) in four plants.

| Life stages | Type of host plant |  |  |  |
| --- | --- | --- | --- | --- |
|  | Mandarin | Curry leaf plant | Himalayan lime | Orange jessamine |
| Egg | 4.16 ± 0.40 | 3.33 ± 0.21 | 5.67 ± 0.42 | 3.67 ± 0.21 |
| Nymph | 5.50 ± 0.34 | 4.83 ± 0.16 | 6.83 ± 0.47 | 5.16 ± 0.47 |
| Adult | 10.67 ± 0.33 | 8.5 ± 0.22 | 9.50 ± 0.42 | 9.33 ± 0.42 |

Additionally, the curry leaf plant demonstrated the fastest rates of oviposition and nymphal development, suggesting that it serves as a more suitable host for the growth and development of the Asian citrus psyllid.

### Survival and death records

The survivability of the Asian citrus psyllid on host plants was assessed across all four treatments (see Table 6). The overall nymphal death rate was highest in the mandarin and Himalayan lime at 41%, while it was lowest when all four host plants were present at 21%. These findings suggest that nymphal survivability increases in the presence of multiple host plants. Conversely, the highest nymphal death rate was observed with the mandarin and Himalayan lime.

**Table 6.** Survival and death records for nymphs of *D. citri* (in percent).

| **Treatment** | **Survival (%)** | **Death (%)** |
| --- | --- | --- |
| Mandarin and Himalayan lime. | 59 | 41 |
| Mandarin and curry leaf plant. | 73 | 27 |
| Mandarin and orange jessamine. | 63 | 37 |
| Mandarin, Himalayan lime, curry leaf plant and orange jessamine. | 79 | 21 |

In an experiment examining four host plants, curry leaf plants exhibited the highest oviposition rate, averaging 22.33 ± 0.33 eggs. In contrast, other choice experiments revealed that orange jessamine had an average oviposition rate of 18.33 ± 0.33 eggs. This finding suggests that the Asian citrus psyllid demonstrates a preference for curry leaf plants when both host plants are available. Furthermore, the highest number of adult psyllids was recorded on the curry leaf plant, indicating that their survivability is optimal on this host.

## Discussions

This experiment confirmed that the Asian citrus psyllid (Diaphorina citri) is capable of breeding and colonizing all four host plants tested. In the choice experiment, D. citri displayed a preference for the orange jessamine plant (Murraya paniculata) for egg-laying. However, when all four host plants were presented together, the psyllids showed a distinct preference for the curry leaf plant (Murraya koenigii) for laying their eggs. This suggests that in the presence of both orange jessamine and curry leaf plants, the psyllids favor the curry leaf plant. Furthermore, adult survival rates were highest on the curry leaf plant, and the overall life cycle of the psyllid was the shortest when this plant was used, lasting only 16.66 days.

The findings on host plant preference between mandarin and Himalayan lime indicated a significant difference (p < .05) in the number of eggs deposited on each plant. Mandarin supported a greater number of eggs (13.67 ± 0.67), nymphs (12.67 ± 1.45), and adults (5.33 ± 0.67), suggesting that Himalayan lime is an inadequate host plant for the Asian citrus psyllid. This outcome stands in contrast to the conclusions drawn by studies [2] and [6], which found that lime was favored by D. citri for oviposition and development. The variation may be attributed to the specific lime species examined in those studies, which differ from the Himalayan lime currently present in Bhutan. Furthermore, a study by [14] established that the Himalayan lime found in Bhutan is a hybrid of Citrus reticulata and Citrus medica (citron).

The results of the comparison between the two host plants, mandarin and curry leaf, reveal a significant difference (p < .05) in the total number of eggs, nymphs, and adults. The curry leaf plant recorded the highest number of eggs (17.67 ± 0.33), while the mandarin plant had only 10.33 ± 0.88. This suggests that Asian citrus psyllids have a preference for curry leaf plants over mandarins for oviposition and breeding. This finding aligns with the study conducted by [17], which concluded that curry leaf plant seedlings are the most susceptible group for the development of all three life stages of D. citri. Consequently, curry leaf plants are employed in research as host plants for D. citri.

The comparison of preferences between the two host plants, mandarin and orange jessamine, reveals a highly significant difference (P < 0.05) in the total counts of eggs, nymphs, and adults. Orange jessamine exhibited higher figures, with an average of 18.33 ± 0.33 eggs, 16.67 ± 0.33 nymphs, and 12.67 ± 0.33 adults, in contrast to mandarin. This finding corroborates the research of [5], which identifies orange jessamine as a preferred host plant for D. citri. Furthermore, the same report indicates that orange jessamine is either resistant or at least tolerant to Huanglongbing disease.

In a study examining four host plants, significant differences (P < .05) were observed in the number of eggs, nymphs, and adults laid by D. citri across the different plants. A Bonferroni multiple comparison test indicated that the curry leaf plant had the highest average number of eggs (22 ± 0.33), while the Himalayan lime exhibited the lowest average (6.67 ± 0.88). Moreover, the curry leaf plant also supported the highest number of adult D. citri (17.67 ± 0.33), whereas no adults were found on the Himalayan lime (0.00 ± 0.57). Although oviposition occurred on all four host plants, higher survival rates were observed on the curry leaf plant and orange jessamine compared to mandarin and Himalayan lime. These findings suggest that the curry leaf plant is the preferred host for oviposition, as well as for nymphal and adult development.

The survivability of the Asian citrus psyllid on various host plants was evaluated across all four treatments (refer to Table 6). The overall nymphal death rate was highest for mandarin and Himalayan lime plants at 41 percent, while it was lowest among the four host plants combined, reaching only 21 percent. These findings suggest that nymphal survivability improves when multiple host plants are available. The elevated nymphal death rate observed in mandarin and Himalayan lime may be due to physiological factors, such as the hardiness and leathery texture of their leaves, as well as the composition and concentration of volatile compounds and odors that might influence the behavior of the Asian citrus psyllid [12,13,16].

In an experiment involving four host plants, the highest rate of oviposition was recorded on the curry leaf plant, with an average of 22.33 ± 0.33 eggs. In contrasting choice experiments, the orange jessamine plant displayed the highest oviposition rate, averaging 18.33 ± 0.33 eggs. This suggests that the Asian citrus psyllid shows a preference for the curry leaf plant over orange jessamine when both host plants are present. Furthermore, the largest number of adult psyllids was found on the curry leaf plant, indicating this plant provides the greatest survivability for the psyllids.

## Conclusions

This experimental study reveals that the curry leaf plant is the preferred host for the Asian citrus psyllid, closely followed by orange jessamine. These two plants have the potential to serve as effective trap crops in mandarin orchards, mitigating the damage caused by this pest. The insights gained from this study are promising and could be implemented in existing citrus-growing regions of Bhutan. As the curry leaf plant is readily available in the wild, it can be utilized as a trap crop to manage Asian citrus psyllids by concentrating them in a limited area within citrus orchards. These trap crops can be removed after infestation by psyllid eggs, prior to the emergence of adult forms. Integrating this approach into current management practices in Bhutan’s citrus orchards could aid in revitalizing the citrus industry and provide a viable alternative to chemical treatments for disease control.

### Recommendations

The study was conducted in a controlled laboratory environment, creating opportunities for further research on the preferences of alternative host plants by Asian citrus psyllids in orchard settings. There are several plants, including grapefruit, citron, and box orange, that have yet to be explored for their effectiveness in attracting Asian citrus psyllids. Additional research on these plants could provide valuable insights into their attractiveness to Asian citrus psyllids in Bhutan.

## Acknowledgments

The authors would like to thank the staff at the National Plant Protection Centre in Simtokha and the Agriculture Research and Development Centre in Bajo for their support and the facilities provided during this study.

## Notes

### Competing Interest Statement

The authors have declared no competing interest.

